# MC1R reduces scarring and rescues stalled healing in a preclinical chronic wound model

**DOI:** 10.1101/2022.11.30.518516

**Authors:** Yonlada Nawilaijaroen, Holly Rocliffe, Shani Austin-Williams, Georgios Krilis, Antonella Pellicoro, Kanheng Zhou, Yubo Ji, Connor A Bain, Alastair M Kilpatrick, Yuhang Chen, Asok Biswas, Michael Crichton, Zhihong Huang, Stuart J Forbes, Andrea Caporali, Jenna L Cash

**Affiliations:** Centre for Inflammation Research, Institute for Regeneration and Repair, 4-5 Little France Drive, University of Edinburgh, Edinburgh, EH16 4UU; Biochemical Pharmacology, William Harvey Research Institute, Barts and the London School of Medicine and Dentistry, Charterhouse Square, London, EC1M 6BQ, UK; Centre for Cardiovascular Research, The Queen’s Medical Research Institute, 47 Little France Crescent, University of Edinburgh, Edinburgh, EH16 4TJ; School of Science and Engineering, Fulton Building, University of Dundee, Dundee, DD1 4HN, UK; Institute of Mechanical, Process and Energy Engineering, School of Engineering and Physical Sciences, Heriot-Watt University, Edinburgh, EH14 4AS, UK; Centre for Regenerative Medicine, Institute for Regeneration and Repair, 4-5 Little France Drive, University of Edinburgh, Edinburgh, EH16 4UU; Department of Pathology, Western General Hospital, Crewe Road, Edinburgh, EH4 2XU, UK

## Abstract

Cutaneous healing results in scarring with significant functional and psychological sequelae, while chronic non-healing wounds represent repair failure often with devastating consequences, including amputation and death. Due to a lack of effective therapies, novel interventions addressing scarring and chronic wounds are urgently needed. Here, we demonstrate that harnessing melanocortin 1 receptor with a selective agonist (MC1R-Ag) confers multifaceted benefits to wound repair. MC1R-Ag accelerates wound closure and re-epithelialization while improving wound bed perfusion and lymphatic drainage by promoting angiogenesis and lymphangiogenesis. Concomitant reductions in oxidative stress, inflammation and scarring were also observed. To evaluate the therapeutic potential of targeting MC1R in pathological healing, we established a novel murine model that recapitulates the hallmarks of human non-healing wounds. This model combines advanced age and locally elevated oxidative stress. Remarkably, topical application of MC1R-Ag restored repair, whereas disrupting MC1R signalling exacerbated the chronic wound phenotype. Our study highlights MC1R agonism as a promising therapeutic approach for scarring and non-healing wound pathologies, and our chronic wound model as a valuable tool for elucidating ulcer development mechanisms.

## INTRODUCTION

Skin is a fascinating and complex barrier organ comprising multiple layers and cell types that are functionally distinct. Repair of adult cutaneous wounds is typically described in terms of four overlapping, sequential phases: coagulation, inflammation, proliferation/migration and remodelling, culminating in scar formation. Most wounds progress through the phases of healing uneventfully, with the magnitude of the inflammatory response linked to the extent of the resultant scar. However, a growing number of wounds fail to progress through the healing phases in an orderly and timely manner, resulting in a persistent open skin ulcer (1,2). These chronic wounds (CWs), including venous leg ulcers (VLU), diabetic foot ulcers (DFU) and pressure ulcers (PU), become ‘stuck’ in a chronically inflamed state, unable to heal effectively. CWs represent a global and escalating health burden due to a sharp rise in worldwide incidence of key comorbidities, including diabetes, obesity and advanced age. CW complications include infection, amputation, sepsis and death; hence they impose substantial morbidity and mortality on the individual, healthcare systems and society, with approximately £12 billion pa spent by the NHS on 3.8 million wounds (3–5).

Whilst many other fields have made inspiring leaps in understanding their respective pathologies, resulting in the development of promising therapeutics, research into CWs has long been neglected. Firstly, access to human wound biopsies is not trivial since the necessary infrastructure to obtain these is not readily available to many researchers. Secondly, the lack of a <u>humane animal model</u> that faithfully <u>recapitulates the key features</u> of human CWs has severely hampered CW research. These issues have therefore compromised innovation in new therapies, clinical practices and biomarkers (6–9).

To remove a major barrier for CW research, we have developed an ethical, reproducible, genetically tractable model that reproduces key features of human CWs, including hyperproliferative epidermis, parakeratosis, spongiosis, vasculitis, fibrinous exudate, erythrocyte extravasation, wound expansion and slough generation. We specifically sought to develop a model that is broadly relevant across the ulcer categories of VLU, DFU and PU, rather than specifically modelling a certain category. Whilst each type of CW is associated with certain comorbidities, for example PU with conditions causing immobility; at their core they share key features. The overwhelming majority of skin ulcers develop in the over 50s, with 95 percent occurring in the over 60s. In addition, VLU, DFU and PU have been shown to exhibit high levels of wound bed oxidative stress. (3,5,10). We therefore used aged animals (18 months) and local elevation of oxidative stress to develop our preclinical model.

This novel model enables the *in vivo* study of candidate therapies to rescue derailed healing responses. Herein, we explore the therapeutic potential of Melanocortin 1 Receptor (MC1R) agonism for treatment of acute and chronic wounds. MC1R is a GPCR expressed peripherally in melanocytes and immune cells, with roles in pigmentation and regulation of anti-oxidative stress mechanisms following UV exposure (11,12). With a functional MC1R, eumelanin, a black pigment, is preferentially produced. A highly polymorphic gene, *MC1R* loss-of-function SNPs result in preferential production of the yellow-orange pigment, phaeomelanin (13). The eumelanin-to-phaeomelanin ratio determines skin and hair colour in humans and coat colour in other mammals. A connection between MC1R and the red-hair phenotype was first established by Valverde *et al* in 1995, however, since then, variants of MC1R have been linked to an increased risk for skin cancer, UV damage, photoaging and more recently with increased severity of post-burn hypertrophic scarring (12–14). Several studies have explored the role of the receptor in inflammatory responses, including animal models of arthritis. These works have demonstrated that delivery of an MC1R agonist can dampen immune cell recruitment and/or activation and thus inflammation, which in the case of the arthritic joint can prevent cartilage degradation (15,16). To date, no studies have explored the role of MC1R in acute and chronic skin wounds, however, Muffley *et al* have described dynamic spatial localisation of MC1R during cutaneous wound repair, including expression by keratinocytes and immune cells in the acute wound bed (11). Here, we demonstrate the multi-faceted mode of action of a selective MC1R agonist (BMS-470539), which promotes wound angiogenesis and lymphangiogenesis whilst accelerating wound closure and reducing scarring. In chronic non-healing wounds, MC1R agonist administration is sufficient to restore effective skin repair.

## RESULTS

### Multifaceted impact of MC1R agonism in acute wound repair

To investigate the role of MC1R in cutaneous wound repair we used a selective MC1R agonist, BMS-470539 (MC1R-Ag), as the endogenous agonist αMSH is also a ligand for MC3R and MC5R. MC1R-Ag or vehicle control were administered topically to 4 mm full-thickness dorsal excisional wounds immediately after injury (Fig. 1A). Macroscopic analysis revealed accelerated wound closure in MC1R-Ag-treated wounds, with maximum differences in wound area noted at 3 and 7 days post-injury (dpi; Vehicle 7 dpi, 16.6 ± 3.8%; MC1R-Ag, 7dpi, 4 ± 2.7% of original area Fig.1B). Key repair parameters were assessed, including epithelial tongue length and scab presence as measures of reepithelialisation. An epithelial tongue is the portion of epidermis that has proliferated and migrated under the scab and was found to be significantly longer at 3 dpi with MC1R-Ag-treatment (Vehicle, 270 ± 19 µm; MC1R-Ag, 399 ± 22 µm Fig.1C). Using optical coherence tomography (OCT) imaging of the dynamic changes in epidermal thickness during wound healing, we demonstrate that harnessing MC1R results in earlier proliferation of wound edge epidermis (peak at 3 dpi vs 5 dpi with vehicle-treatment; Fig. 1D), this was followed by a quicker reduction in epidermal thickness by 7 dpi (Vehicle, 140 ± 8 µm; MC1R-Ag, 93 ± 9 µm, Fig. 1D, E). Since re-epithelialisation was completed earlier in MC1R-Ag-treated wounds, this resulted in reduced scab presence at 7 dpi (Vehicle, 71 ± 8.5%; MC1R-Ag, 18 ± 7% of wounds; Fig. 1F,G). We utilised an *in vitro* scratch wound assay to determine whether the observed effects of MC1R-Ag on re-epithelialisation could be through direct actions on epidermal keratinocytes. Indeed, MC1R-Ag accelerated keratinocyte scratch wound closure (Fig.S1A-C).

**Figure 1.**
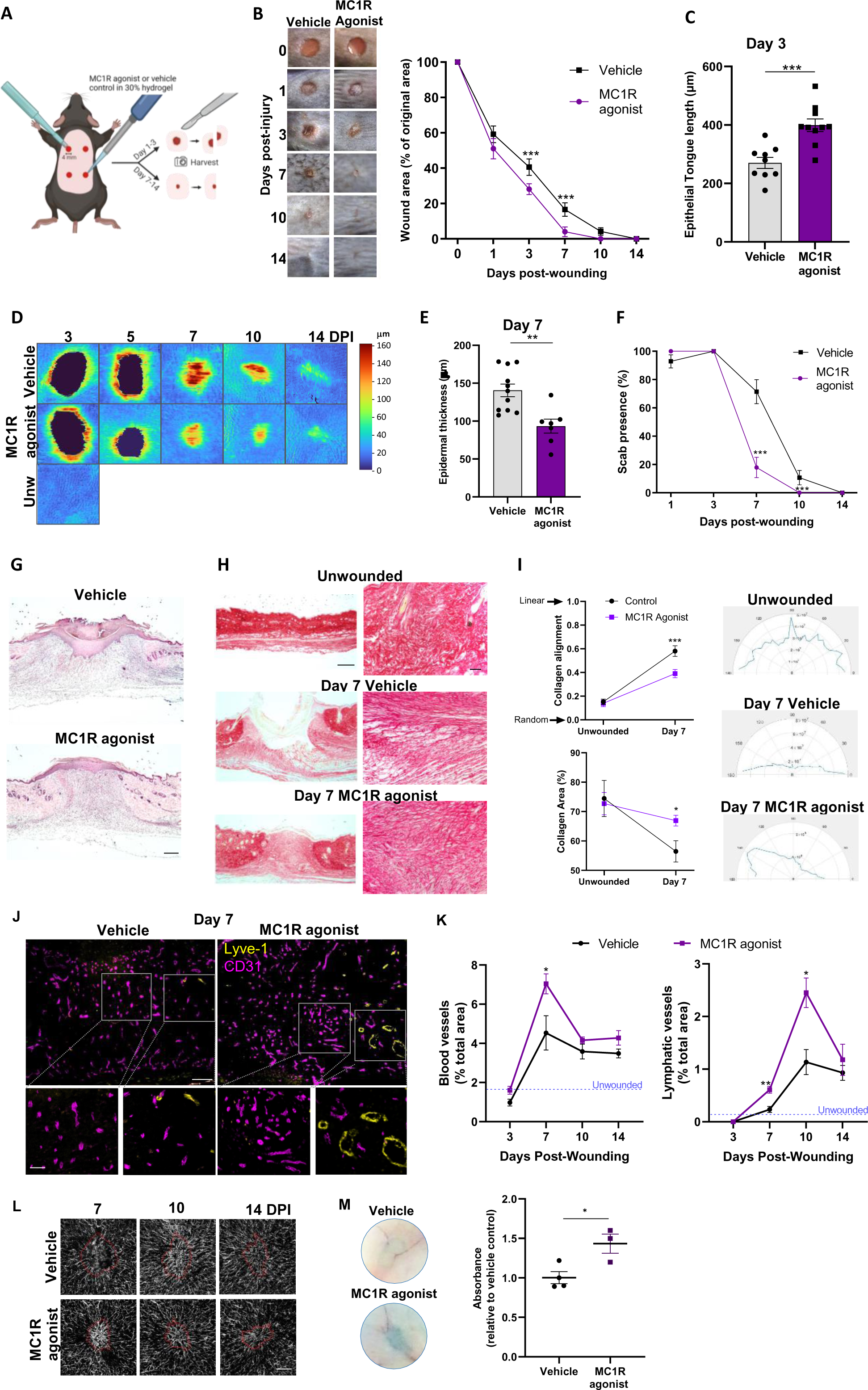
Multifaceted impact of MC1R agonism on acute wound repair. Four 4 mm excisional wounds were made to the dorsal skin of 8 week old C57Bl/6J mice. Vehicle (PBS) or MC1R agonist (BMS-470539) in 30% Pluronic hydrogel was administered topically into the wound immediately after wounding. (**A)** Schematic diagram illustrating the location and processing of acute skin wounds. **(B)** Representative macroscopic photos and quantification of wound areas 1-14 dpi. n = 7 biological replicates for each group from three independent experiments. **(C)** Epithelial tongue length at 3 dpi. 9 wounds (vehicle), 10 wounds (MC1R agonist). **(D)** Heatmap generated from Optical coherence tomography (OCT)-derived images illustrating epidermal thickness immediately around the wound at 3 and 5 dpi, and in the wound bed at 7, 10 and 14 dpi. 6 wounds per group. **(E)** Epidermal thickness at 7 dpi, with 11 (vehicle) and 7 (MC1R agonist) wounds. **(F)** Scab presence 1-14 dpi. n = 7 biological replicates for each group from three independent experiments. **(G)** Representative H&E-stained wound mid-sections at 7 dpi illustrating scab loss and smaller wound area in MC1R-agonist treated wounds (5x, lpf). **(H)** Picrosirius red-stained collagen in unwounded skin and wound mid-sections at 7 dpi. Lpf (5x, left), hpf (20x, right). **(I)** Collagen alignment and area in unwounded skin and 7 dpi wounds, with 11 (vehicle) and 7 (MC1R-Ag) wounds per group. Right: Polar plots illustrating collagen fiber alignment. **(J)** Representative CD31 (magenta) and Lyve-1 (yellow) immunostaining of vehicle- and MC1R agonist-treated wounds at 7 dpi (vehicle, n=6; MC1R agonist, n=5 per treatment group from three independent experiments). **(K)** Quantification of CD31^+^ (blood vessel) area and Lyve-1^+^ (lymphatic vessel) area per hpf in wounds 3-14 dpi. Area in unwounded skin denoted by dotted blue line. **(L)** OCT angiography of blood flow demonstrating increased perfusion of MC1R agonist-treated wounds at 7dpi. Red line demarcates wound edge identified using OCT structural image. **(M)** Representative macroscopic photos and quantification of skin-draining lymph nodes of mice 10 dpi, where Evans blue was injected into the wound bed to assess lymphatic drainage capacity of wounds treated with vehicle control or MC1R agonist at 0 dpi. Data are expressed as mean ± SEM. * p<0.05, ** p<0.01, *** p<0.001 by two-way ANOVA with Bonferroni’s multiple comparison test (B, F, I, K) or Student’s unpaired t test (C, E, M) relative to vehicle-treated wounds. Dpi; days post-injury.

Physiological skin repair culminates in formation of a scar, which predominantly consists of linear bundles of collagen fibers, as opposed to normal dermis, which exhibits a basketweave appearance (1,17). To quantify scarring, we measured collagen fiber orientation using a novel app, FIBRAL and picrosirius red-stained sections. Unwounded skin exhibited abundant collagen with randomly-oriented fibers corresponding with a low alignment index (0.152 ± 0.024; Fig. 1H-I). In vehicle-treated day 7 wounds, collagen was laid down in a more linear fashion with a high alignment value, typical of a scar. Wound treatment with MC1R-Ag, however, resulted in collagen deposition with a more random orientation closer to unwounded dermal collagen organisation, which equated to reduced alignment in comparison to vehicle-treated wounds (0.39 ± 0.036 MC1R-Ag vs 0.58 ± 0.045 vehicle; Fig. 1H-I). We also assessed collagen quantity, finding that MC1R-Ag accelerated the rate of collagen deposition (66.9 ± 1.85% MC1R-Ag vs 56.5.2 ± 3.64% vehicle vs unwounded 74.5 ± 6.17%; Fig. 1H-I). Thus a downstream consequence of wound MC1R-Ag treatment is reduced scarring through improved collagen deposition quality and quantity.

### Harnessing MC1R drives angiogenesis and lymphangiogenesis at sites of cutaneous repair

Angiogenesis and lymphangiogenesis are processes that principally occur either during early development or during wound repair or inflammation (18). Wound angiogenesis is required to enable delivery of oxygen and nutrients to the metabolically demanding repairing tissue, with impaired angiogenesis observed in aberrant healing scenarios including DFUs (5,19). Lymphatic vessels are thought to be important in wound healing as they drain interstitial fluid to reduce tissue swelling and permit the transportation of immune cells to lymph nodes (20). We therefore investigated the effect of MC1R-Ag on angiogenic and lymphangiogenic responses during wound repair (Fig. 1J-M). As we and others have previously shown (21–23), the time course of angiogenesis preceded that of lymphangiogenesis with angiogenic sprouting occurring as early as 3 dpi, peaking at 7 dpi and vascular remodelling, including trimming of superfluous vessels occurring thereafter (Fig. 1J,K). The lymphangiogenic response, however, was evident at 7 dpi, peaking at 10 dpi (Fig. 1J,K). Wound treatment with MC1R-Ag profoundly enhances the angiogenic (155% at 7 dpi) and lymphangiogenic (216% at 10 dpi) responses with increased vessel density over vehicle-treated wounds (Fig. 1J,K). To validate that the CD31+ cell areas identified through immunofluorescence staining represented functional and perfused blood vessels, we performed optical coherence tomography (OCT) angiography. Notably, OCT angiography revealed substantial wound bed perfusion at 7 days post-injury in the MC1R-treated group, which subsequently declined over time consistent with vessel remodelling (Fig. 1L). To investigate whether the enhanced wound vascularization observed with MC1R-Ag treatment was mediated through direct effects on endothelial cells, we employed a microvascular endothelial cell tube formation assay. MC1R-Ag significantly increased readouts of *in vitro* angiogenesis, including total mesh area, branching interval and total segment area (Fig. S1D-F). MC1R siRNA knockdown demonstrated that these angiogenic effects of BMS-470539 are mediated by MC1R (Fig. S1F). To assess wound lymphatic vasculature function, we employed a lymphatic drainage assay, whereby Evans Blue dye injected into wounds was quantified in skin-draining lymph nodes. We confirm that the observed increase in lymphatic vessel area translated to improved lymphatic drainage in MC1R-Ag-treated wounds (Fig. 1M). Collectively, we demonstrate that BMS-470539 acts via endothelial cell MC1R to drive angiogenesis and lymphangiogenesis, resulting in improved wound bed blood supply and lymphatic drainage.

### MC1R modulates the wound microenvironment

We monitored leukocyte recruitment dynamics during dermal wound repair, finding that MC1R-Ag administration modeslty inhibits neutrophil recruitment to excisional cutaneous wounds (Fig. S2A). Surprisingly, MC1R-Ag increased F4/80^+^ MΦ recruitment to day 7 wounds (Fig. S2B), however we noted that MΦ distribution in the skin adjacent to the wound was limited by MC1R-Ag treatment, unlike vehicle-treated day 7 wounds where an F4/80^+^ infiltrate is present in the wound adjacent dermis (Fig. S2C-D). We noted no clear change in wound MΦ morphology, however, a marked improvement in erythrocyte clearance (Prussian blue^+^ cells; Fig. S2E) was observed, along with significantly reduced wound TNFα (Fig. S2F) and oxidative stress (dihydrorhodamine^+^ cells; Fig. S2G). Collectively, these data suggest that MC1R agonism modulates the wound microenvironment, possibly resulting in a shift towards the repair phase (proliferation/migration and remodelling).

### Lack of functional MC1R impairs wound healing

In humans and mice, lack of a functional MC1R due to spontaneous mutation, results in the red-hair phenotype. To establish the impact of MC1R on wound healing, we generated 4 mm wounds in MC1Re/e mice, which harbour a non-functional receptor (ginger hair; C57Bl/6J background) and in wildtype (black hair; C57Bl/6J) mice. We discover that lack of functional MC1R profoundly impaired healing, with significantly delayed wound closure detectable at 1, 7, 10 and 14 dpi (7 dpi – MC1Re/e, 27.4 ± 2.1%; C57B/6J, 7.36 ± 0.83% of initial wound area; Fig. 2A-D). Incomplete reepithelialisation was evident in MC1Re/e wounds 7 dpi with presence of epithelial tongues and gap between them, in comparison to reformation of an intact epidermal layer in C57Bl/6J control wounds (Fig. 2D). This resulted in delayed scab loss (an indicator of complete reepithelialisation), with 95% of MC1Re/e wounds retaining a scab at 7 dpi in comparison to 68.8% of wildtype control wounds (MC1Re/e + vehicle, 95 ± 5.1%; MC1Re/e + BMS-470539, 90.2 ± 6.1%; C57Bl/6J, 68.8 ± 7.84; Fig. 2E). Furthermore, abundant NETs were observed in MC1Re/e wounds suggesting a role for endogenous MC1R ligands in regulating their formation (Fig. 2F). Importantly, MC1R agonist, BMS-470539, is unable to impact the wound healing response in MC1Re/e mice (Fig. 2A,B,E). Collectively, these data establish that MC1R is required for optimal healing.

**Figure 2.**
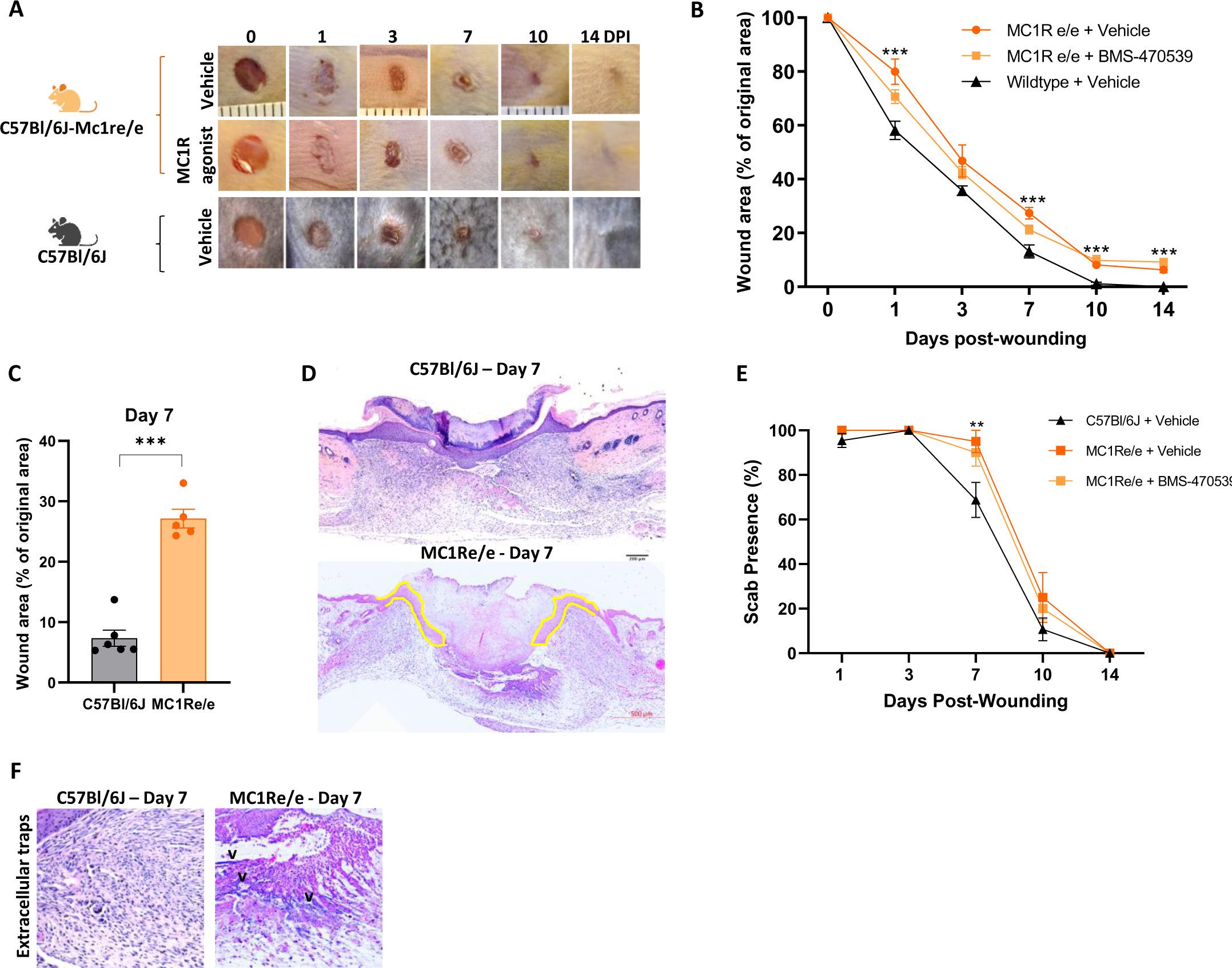
Lack of functional MC1R impairs acute wound healing. Four 4 mm excisional wounds were made to the dorsal skin of MC1Re/e and C57Bl/6J mice. Vehicle (PBS) or MC1R agonist (BMS-470539) in 30% Pluronic hydrogel was administered topically into the wound immediately after wounding. **(A)** Representative macroscopic photos of vehicle- and BMS-470539-treated wounds in MC1Re/e mice in comparison to C57Bl/6J vehicle-treated wounds (also see Fig. 1B). **(B)** Longitudinal macroscopic quantification of wound areas in MC1Re/e and Wildtype mice at 1-14 dpi. 5-7 biological replicates for each group from 2-3 independent experiments. **(C)** Wound area at 7 dpi in C57Bl/6J (n=6) and MC1Re/e mice (n=5). **(D)** Representative H&E-stained wound mid-sections at 7 dpi in MC1Re/e and wildtype mice. Epithelial tongues highlighted in yellow, illustrating incomplete reepithelialisation in MC1Re/e mouse wounds. **(E)** Scab presence 1-14 dpi in MC1Re/e and wildtype mice. 5-7 biological replicates for each group from three independent experiments. **(F)** Extracellular traps (likely NETs, marked with **v**) visible in H&E-stained MC1Re/e wound mid-sections but not wildtype controls 7 dpi. Data are expressed as mean ± SEM. ** p<0.01, *** p<0.001 by two-way ANOVA with Bonferroni’s multiple comparison test (B,E) or Student’s unpaired t test (C) relative to C57Bl/6J wounds. 5 MC1Re/e + vehicle, 5 MC1Re/e + BMS-470539 and 8 C57Bl/6J + vehicle-treated animals per group from 2 independent experiments.

Since acute wounds typically heal without complications, we asked whether MC1R agonism would be of benefit in the context of non-healing wounds.

### Chronic wound model reproduces hallmarks of the human pathology

Given that the majority of acute wounds heal without significant complications, and CWs represent a profound and escalating burden to individuals and healthcare systems (3), we sought to develop a preclinical CW model to act as a platform for testing of novel interventions such as MC1R-Ag. Equally importantly, the provision of a humane CW model has long been recognised as essential to better understand the disease process without the risk of clinical harm (7). Here, we utilised the mouse as our model organism, since they are amenable to genetic manipulation, reproduce rapidly and are easy to house and maintain.

As described by Frykberg and Banks, CWs have variable underlying pathologies, including diabetes, immobility and venous stasis and can be divided into categories based on these, such as DFU, PU and VLU (5). We deliberately sought to develop a model that is broadly relevant across the ulcer categories, rather than specifically modelling a certain category. Whilst each type of CW is associated with certain comorbidities, at their core they share key features. The overwhelming majority of skin ulcers develop in the over 50s, with 95 percent occurring in the over 60s. In addition, VLU, DFU and PU have been shown to exhibit high levels of wound bed oxidative stress (3,5,10). Taking inspiration from the advantages and limitations of current murine models (6,7,25,26), we developed our model by combining advanced age and local elevation of oxidative stress (Fig. 3A).

**Figure 3.**
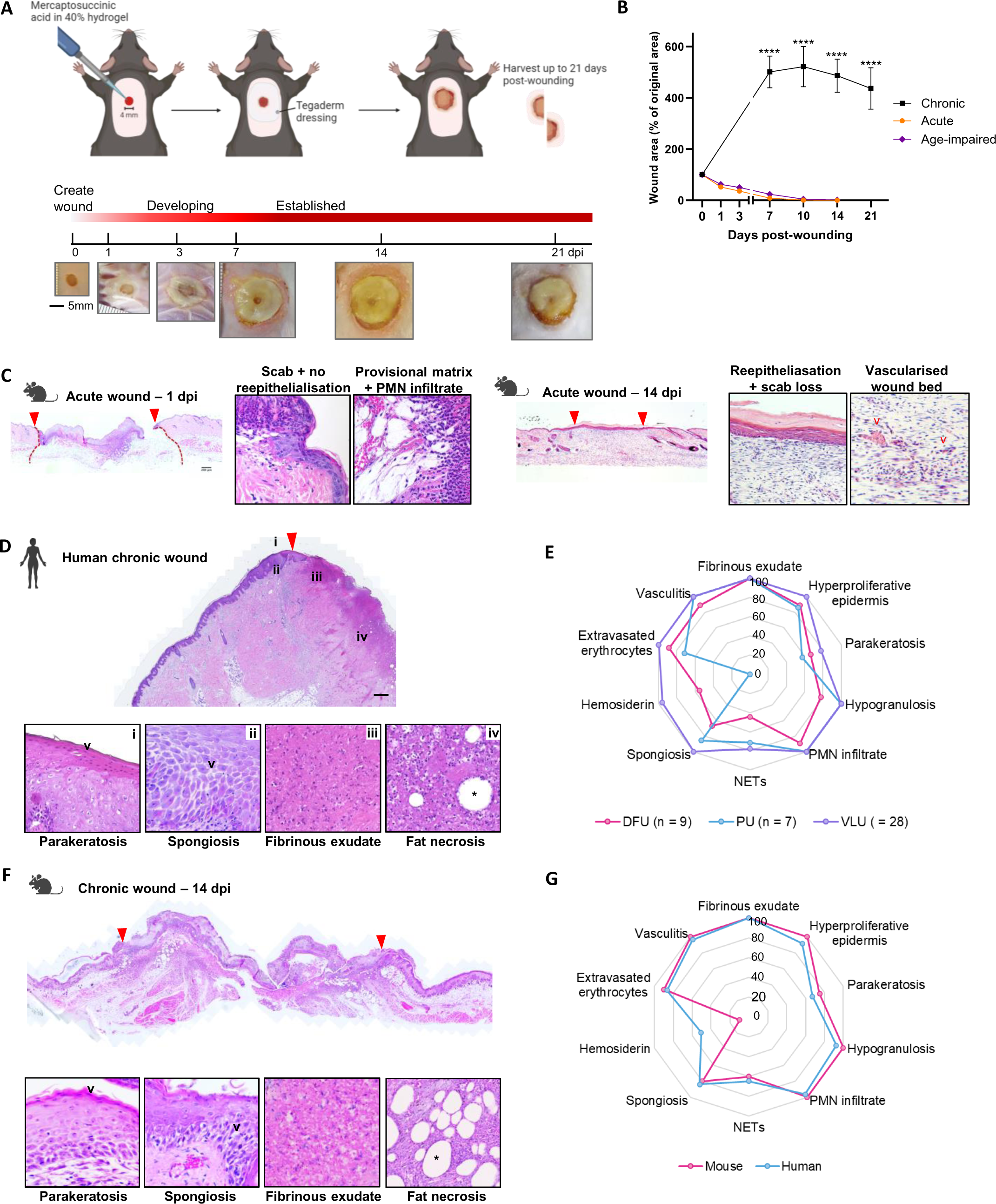
Established a humane chronic wound model that recapitulates key features of human non-healing wounds. **(A)** Top - Schematic detailing the method used to create chronic wounds (CWs) in aged animals. A single 4 mm excisional wound was made to the upper dorsal skin of 18-22 m old C57Bl/6J mice. Mercaptosuccinic acid suspended in 40% Pluronic hydrogel was topically administered to the wound immediately after injury, followed by placement of a Tegaderm wound dressing and wound harvest up to 21 dpi. Bottom – Macroscopic photos showing CW development from 0–21 dpi. **(B)** Longitudinal macroscopic quantification of wound areas 1-14 dpi in acute (young 8 w old mice; n=8) and age-impaired (18-24 m old mice; n=8) healing versus 7-21 dpi chronic non-healing wounds (n=15 at 14 dpi, n=8 at 7, 10 and 21 dpi). Data expressed as mean ± SEM, **** p<0.0001. **(C)** Representative low power field (lpf) **mouse acute wound** H&E sections at 1 dpi (left) and 14 dpi (right) with key histological features shown below in high power field (hpf). n=8 biological replicates. **(D)** Representative lpf **human CW** H&E section (n=44 patients) with key histological features shown below in hpf. **(E)** Radar plot indicating mean prevalence (%) of key CW features in human DFU (n=9), VLU (n=28) and PU (n=7). **(F)** Representative lpf **mouse CW** H&E sections at 14 dpi with key histological features shown below in hpf. n=10 biological replicates from 3 independent experiments. **(G)** Radar plot indicating mean prevalence (%) of key features observed in human CWs (DFU, VLU and PU, n=44), in comparison to the murine preclinical CW model (n=10). Red arrow heads indicate wound margins, * indicates fat necrosis. Dpi, days post-injury.

A 4 mm excisional wound was made to the dorsal skin of aged mice, topical Glutathione peroxidase (GPx) inhibitor, mercaptosuccinic acid (MSA) was applied, following by a transparent Tegaderm dressing (Fig. 3A). The wounds expanded into the surrounding tissue 5-fold over the first 7 dpi and stabilised at this size for the duration of the experiment (Fig. 3A-B). The tissue initially took on a macerated (white) appearance (approx. 1 dpi), followed by a thin red border appearing around the macerated area at 3 dpi but absent by 7 dpi. The wounds produced exudate and slough (as seen in human CWs), and were well tolerated by the animals (Fig. 3B and S3). Wound architecture was compared between the well established acute excisional wound model (Fig. 3C-D), the new CW model (Fig. 3F-G) and a cohort of 44 human CWs (28 VLU, 9 DFU, 7 PU; Fig. 3D-E, Fig. S4). During successful skin repair, at only 1 dpi a scab has formed as a result of haemostasis, reepithelialisation is not evident, provisional matrix has been deposited and a rich PMN infiltrate is notable (Fig. 3C). In contrast, at 14 dpi, reepithelialisation is complete, scab loss has occurred, few PMN remain and the wound bed is well vascularised (Fig. 3C). In human CWs we find that fibrinous exudate, hyperproliferative epidermis, PMN infiltrate and vasculitis are consistently present, being identified in 86-100% of DFU, VLU and PU (Fig. 3D-E). Hypogranulosis (78-100%), extravasated erythrocytes (71-100%), parakeratosis (57-78%) and spongiosis (67-100%) were also common histomorphological features (Fig. 3D-E). Hemosiderin deposition proved to be the most variable feature, found in all VLU, 56% of DFU but none observable in PU. This likely relates to specific co-morbidities including venous stasis that are most commonly found in patients with VLUs (Fig. 3D-E). In our preclinical CW model, macroscopic and histomorphological analysis revealed excellent recapitulation of hallmarks of human CWs (Fig. 3F-G). Notable, were a severe lack of granulation tissue formation, substantial and persistent fibrinous exudate, presence of extravasated erythrocytes and subcutaneous fat necrosis. Major disturbance to epidermal physiology was evident through hyperproliferation, spongiosis and parakeratosis, along with a complete failure of wound reepithelialisation. Whilst exudate and slough were abundant, scab formation was absent (Fig. 3F-G).

Having established a humane CW model that accurately mimics many of the hallmarks of human CWs, we then sought to evaluate whether MC1R-Ag administration could improve healing outcomes, thus representing a potential therapeutic approach.

### MC1R agonist restores healing in a preclinical chronic wound model

In the clinic, non-healing wounds are typically managed in the community, with dressing changes and basic wound debridement performed as part of standard wound care (27). Effective wound debridement is thought to partially stimulate healing, but in complex wounds, is typically insufficient to result in complete repair. To assess a potential therapeutic intervention, we therefore debrided the wounds 7 dpi (Fig. 4A,B). Vehicle-control or MC1R-Ag were administered topically in a hydrogel immediately after debridement and at each subsequent dressing change 3 and 5 days later (Fig. 4A,B). Vehicle-treated debrided wounds mimicked the hallmarks of human CWs, including extravasated erythrocytes, PMN infiltrate, fibrinous exudate, spongiosis, parakeratosis, a lack of wound vascularisation and absence of reepithelialisation (Fig. 4B,C), whilst exhibiting a partial stimulation to healing, evident as a 30% reduction in wound area at 14 dpi. However, MC1R-Ag treatment following debridement rescued the healing response, with an additional 33% reduction in wound area over debridement alone by 14 dpi, rising to 68% at 21 dpi. Here, we observed partial reepithelialisation by 14 dpi, as evidenced by formation of epithelial tongues, with complete reepithelialisation at 21 dpi. In addition, minimal fibrinous exudate was present and a vascularised granulation tissue formed (Fig. 4B,D). In contrast, MC1Re/e CWs produced copious exudate both pre and post-wound debridement (Fig. 4E,H) such that 88% exhibited notable exudate 14 dpi in comparison to 56% of vehicle-treated wildtype wounds. Presence of wound exudate was reduced with MC1R-agonist treatment with only 20% of wounds displaying visible exudate 14 dpi, reducing to 0% by 21 dpi (Fig. 4D,H). Importantly, NETosis, spongiosis and hemosiderin deposition were notable features of MC1Re/e CWs, but extravasated erythrocytes were not observed (Fig. 4E,I).

**Figure 4.**
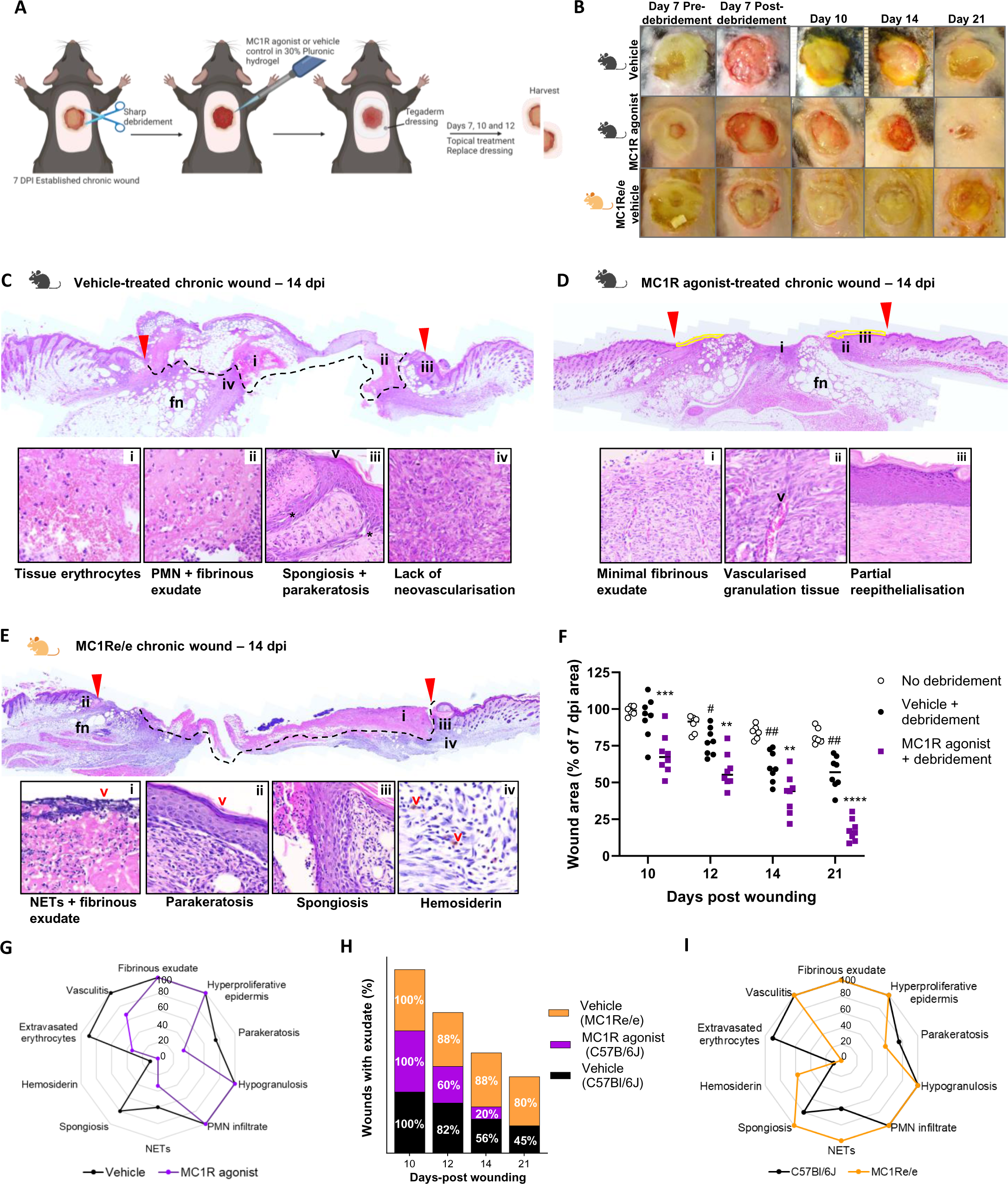
MC1R agonist restores healing in murine chronic wounds. **(A)** Schematic diagram detailing the CW protocol from wound debridement through to harvest. Wounds were debrided of devitalised tissue 7 dpi, with dressing changes and topical delivery of vehicle control or MC1R agonist in 30% Pluronic hydrogel at 7, 10 and 12 dpi and wound harvest up to 21 dpi. **(B)** Representative macroscopic photos showing CWs immediately before and after debridement at 7 dpi, then at 10, 12, 14 and 21 dpi with topical vehicle- or MC1R agonist-treatment. MC1Re/e mice received identical treatment to vehicle-treated C57Bl/6J animals. **(C-E)** Representative lpf H&E-stained wound mid-sections at 14 dpi with key histological features shown below in hpf. **(C)** Vehicle-treated CW in C57Bl/6J mouse, **(D)** MC1R agonist-treated CW in C57Bl/6J mouse. Yellow lines mark epithelial tongues. **(E)** Vehicle-treated CW in MC1Re/e mouse. Red arrow heads mark wound edges, tissue above the black dashed lines is fibrinous exudate (fe). fn, fat necrosis. **(F)** Quantification of wound areas at 10 - 21 dpi, normalised to 7 dpi areas. ** p<0.01 relative to Vehicle + debridement. # p<0.05 relative to No debridement group. **(G)** Radar plot indicating mean prevalence (%) of key histomorphological features in vehicle and MC1R agonist-treated CWs. **(H)** Stacked bar graph indicating prevalence of CWs with visible wound exudate. **(I)** Radar plot indicating mean prevalence (%) of key histomorphological features in C57Bl/6J and MC1Re/e CWs. C57Bl/6J CWs with no debridement (n=6), with debridement and vehicle-treatment (n=7), with debridement and MC1R agonist-treatment (n=5). MC1Re/e CWs with no debridement (n=7) from three independent experiments.

### Dysregulation of POMC-MC1R axis in non-healing DFU

To gain further insight into the mechanism of action of MC1R agonist, we examined expression of *MC1R* and it’s endogenous ligand precursor, pro-opiomelanocortin (POMC). POMC is converted to the endogenous MC1R ligand αMSH through a series of enzymatic cleavages, thus necessitating our assessment of *POMC* expression. We found *POMC* to be broadly expressed in multiple cell types, including vascular endothelial cells, basal and differentiated keratinocytes and highest in fibroblasts and smooth muscle cells (Fig. 5A). Low levels of *POMC* expression were also noted in macrophages, plasma and T cells. *MC1R* was found to be expressed in vascular endothelial cells, melanocytes, fibroblasts and smooth muscle cells, with lower levels detected in keratinocytes (Fig. 5A). We performed CellChat analysis (Jin et al 2021) to assess POMC-MC1R pathway signalling. Interestingly, we found a reduction in interaction strength, but not the number of interactions, was associated with non-healing DFU, but not healing DFU or healthy skin (Fig. 5B). Cell-cell communication network inference revealed extensive cell-cell communication via the *POMC*-*MC1R* pathway in skin and DFU. In healthy skin, the dominant senders are differentiated keratinocytes, basal keratinocytes and plasma cells, with melanocytes the dominant receivers. M2 macrophages, sweat glands, fibroblasts and smooth muscle cells (SMC) act as key mediators, whilst most cell types studied act as influencers except the vascular and lymphatic endothelium and B lymphocytes (Fig. 5C-D). In healing DFU, the lymphatic endothelial cells become weak senders, along with sweat glands. Plasma cells are no longer contributing in any role. SMC, fibroblasts, differentiated and basal keratinocytes and sweat glands become receivers, largely consistent with the role of MC1R in modulating wound contraction, scar deposition, reepithelialisation and revascularisation. Differentiated and basal keratinocytes become mediators and endothelial cells become influencers suggesting their role as gatekeepers of cell-cell communication in the MC1R pathway during healing. In contrast, non-healing DFU largely fail to induce the changes in POMC-MC1R communication observed in healing DFU. MC1R staining of DFU confirmed epidermal and vascular expression (Fig. 5E).

**Figure 5.**
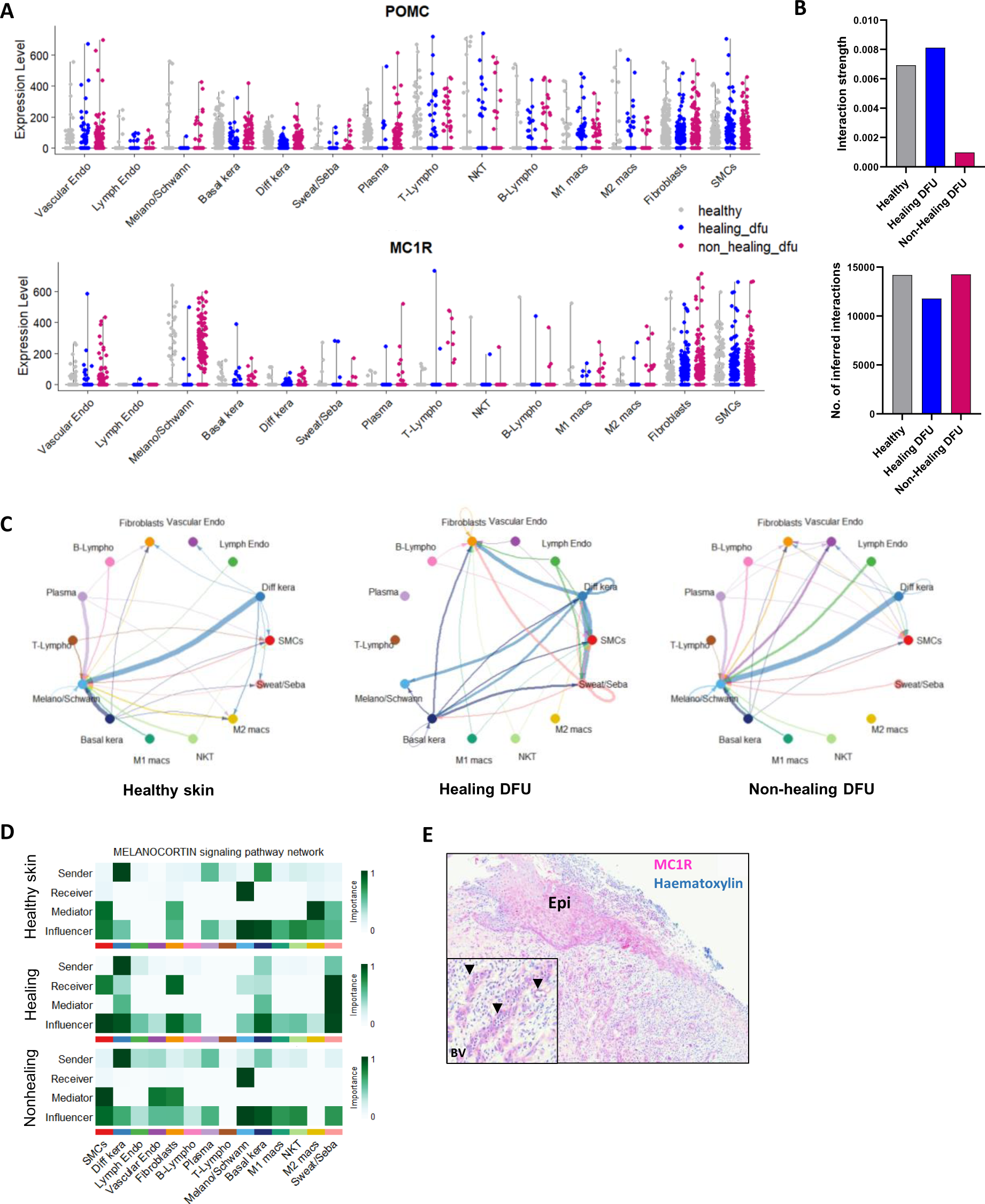
Dysregulated POMC-MC1R axis in human chronic wounds. **(A)** Violin plots depicting POMC and MC1R expression across cell types in healthy skin, healing and non-healing DFU. **(B)** Comparison of the total number of interactions and interaction strength for the POMC-MC1R ligand-receptor pair, illustrating reduced interaction strength in non-healing DFU in comparison to healthy skin and healing DFU. **(C)** Circle plot of inferred POMC-MC1R signalling networks in healthy human skin, healing diabetic foot ulcers (DFU) and non-healing DFU. Edge weight is proportional to the inferred interaction strength. Thicker edge line indicates a stronger signal. Edges colours are consistent with the signalling source. Arrows indicate signal direction **(D)** Heatmap showing the relative importance of each cell group based on the computed four network centrality measures of the MC1R signaling network. **(E)** Representative MC1R (Vector red)-stained DFU demonstrating epidermal and vascular expression. Epi; epidermis. Inset with arrows marking blood vessels, BV. SMC, smooth muscle cells; Diff kera, differentiated keratinocytes; Lymph endo, lymphatic endothelial cells; Melano/Schwann, melanocytes/Schwann cells; NKT, natural killer T cells; Sweat/Seba, sweat/sebaceous glands. **(A-D)** derived from reanalysis of GSE165816 (36).

## DISCUSSION

In this study, we investigated the therapeutic potential of harnessing MC1R in wound healing by employing a selective agonist, BMS-470539. Our findings demonstrate that MC1R agonism accelerated wound closure and reepithelialisation in wildtype but not MC1Re/e mice, which harbour a non-functional receptor. Notably, MC1R agonism improved wound perfusion and lymphatic drainage by promoting angiogenesis and lymphangiogenesis, reducing local oxidative stress and inflammation with a knock-on effect of reduced scarring. While the cellular and molecular mechanisms underlying skin repair, scarring and particularly the development and persistence of CWs remain incompletely understood, our study suggests that targeting MC1R represents a promising strategy to rescue failed healing and mitigate excessive scarring. This therapeutic potential, likely stems from MC1R agonism enabling a more rapid transition to the proliferative phase of wound healing to effect efficient healing. It is worthwhile noting that in the context of drug development, a multi-modal approach targeting diverse aspects of the intricate healing process is generally considered advantageous over single mode-of-action therapies. By simultaneously engaging several mechanisms, drugs with multiple modes-of-action can provide improved efficacy, reduced likelihood of treatment resistance and a more comprehensive modulation of the complex cellular and molecular pathways involved in wound repair.

Given the reported inhibitory effects of BMS-470539 on leukocyte recruitment in other systems (15), we were surprised to observe minimal impact on leukocyte recruitment, however, it is entirely possible that some of the impact on wound healing may be through modulating leukocyte phenotype or via leukocyte lineages out with the scope of this study. Nevertheless, MC1R-Ag acts directly on endothelial cells to drive microvascular endothelial cell sprouting and mesh formation and may therefore hold promise for pathologies characterised by poor vascularisation. MC1R activation also has potential anti-fibrotic properties. Collagen alignment plays a significant role in scar quality, with a disordered collagen architecture being characteristic of an unwounded dermis. Using simulations and multi-scale mathematical modelling, McDougall *et al* derived equations to elucidate the molecular cause of alignment during collagen deposition. Their results suggest that a localised chemoattractant gradient, with greater effects on only those fibroblasts in the vicinity of the wound, produces a collagen pattern that is highly disordered (24). This could be the driver behind reduced collagen alignment in MC1R agonist-treated wounds; macrophages were more restricted to the wound site and could thus provide localised chemoattractants.

To remove a major barrier for CW research, we describe herein the development of an ethical, reproducible and genetically tractable model that reproduces key features of human CWs. We consciously avoided developing a diabetic CW model due to the need for broad applicability and limitations of monogenic murine diabetes models not replicating the polygenic and multifactorial nature of human type 2 diabetes. Inevitably, every preclinical model has limitations, and it’s crucial to acknowledge them, especially when translating findings to the clinic. Regarding our preclinical CW model, key considerations involve the unavoidable anatomical differences between mouse and human skin. While in humans, the epidermis and dermis are thicker, and the skin firmly adheres to the fascia, in mice, the skin is typically not adherent to the underlying fascia, although chronic inflammation can induce adherence in our model (28)(29). Additionally, the duration of human CWs poses a challenge, as modelling years-long persistence is unethical and infeasible in mice with shorter lifespans. Instead, we model CW development and shorter-term persistence, choosing time points that provide the opportunity to directly compare the early repair derailment process to acute wound repair. We anticipate that our preclinical CW model will become a valuable tool to interrogate ulcer formation and persistence along with testing of novel interventions.

## MATERIALS AND METHODS

### BMS-470539

MC1R agonist, BMS-470539, was synthesised by Mimotopes (Minneapolis, USA), reconstituted (1 mM in PBS + 0.1% BSA) and stored at -80°C for up to 6 months. BMS-470539 was diluted from fresh aliquots for each experiment.

### Animals

All experiments were conducted with approval from the University of Edinburgh Local Ethical Review Committee and in accordance with the UK Home Office regulations (Guidance on the Operation of Animals, Scientific Procedures Act, 1986) under PPL PD3147DAB. C57Bl/6JCrl and MC1Re/e mice on a C57Bl/6JCrl background were bred in-house (LFR at BioQuarter site), maintained in conventional cages on a 12:12 light:dark cycle with *ad libitum* access to standard chow and water under a SPF environment. Animals were housed 3-5 per cage in a temperature (22-24C) and humidity controlled room. All animals were housed in the same room throughout their lifespan. Environmental enrichment was provided in the form of dome homes, a tunnel and wooden chew sticks. Health checks were performed on all animals prior to wounding, including baseline weight measurements, malocclusion and skin injuries due to fighting. Animals with malocclusion were excluded from the study and those with small skin injuries due to historic fighting were included providing wounds could be made without involvement of the previous injury area. Only healthy animals that were not involved in previous procedures were used for experiments. The chronic wound model described below was developed following the 3Rs framework (30) and with regular input from the NVS (Named Veterinary Surgeon, Nacho Vinuela-Fernandez). Both the acute and chronic wound model are categorized as moderate severity protocols. No deaths occur during these protocols and weight loss is less than 10% of starting weight (Fig. S6).

### Aging mouse colony

Once animals reached 12 months of age, several parameters of health were monitored on a weekly basis to determine animal welfare. Parameters were weight (as compared to baseline 12 mo weight), body condition scores (as determined by visual and physical examination of each animal in accordance with the University of Edinburgh Veterinary Scientific Service guidelines) and behaviour. Weight loss of 10% and/or an increase in body condition score or evident lack of expected behaviour (interaction, movement) resulted in the animal being assessed by a vet.

### Acute skin wounds

Male and female mice (7-9 weeks old) were randomly assigned a treatment group and anaesthetized with isoflurane (Zoetis, Leatherhead, UK) by inhalation. Buprenorphine analgesia (0.05 mg/kg, s.c, Vetergesic, Amsterdam) was provided immediately prior to wounding and dorsal hair was removed using a Wahl trimmer. Four full-thickness excisional wounds were made to the shaved dorsal skin using sterile, single use 4 mm punch biopsy tools (Selles Medical, Hull, UK). Wounds were photographed with a Sony DSC-WX350 and a ruler immediately after wounding and also on days 1, 3, 7, 10 and 14 post-wounding. Vehicle (PBS) or BMS-470539 were delivered topically by pipette into the wound cavity immediately after wounding (40 μl in a 30% Pluronic F-127 gel (liquid at 4°C, but solidifies at room temperature; Sigma Aldrich). Wound areas were calculated using ImageJ software.

Mice were housed with their previous cage mates in a 28C warm box (Scanbur, Denmark) overnight following wounding, with paper towels used as bedding to avoid sawdust entering the open wounds. Dome home entrances were enlarged to prevent animals scraping their dorsal skin wounds. Animals were moved into clean conventional cages at 22-24C the following morning.

### Chronic skin wounds

Mice (18-22 months, mean weight ± sd; 34.8 g ± 6.2 g) were randomly assigned a treatment group, anaesthetized with isofluorane and placed on a heated mat to maintain body temperature. Buprenorphine analgesia (0.05 mg/kg, s.c, Vetergesic, Amsterdam) was provided immediately prior to wounding and dorsal hair was removed using a BaByliss trimmer. Nair cream (Church and Dwight, Folkestone, UK) was used to remove the remnants of the dorsal hair to create a surface that the Tegaderm dressing would adhere to. Depilatory cream was carefully removed using PBS and tissue paper to avoid chemical irritation. Claws were trimmed to minimise the potential for scratch-induced skin trauma during the protocol. A single 4 mm excisional wound (Selles Medical, UK) was made to the dorsal skin near the shoulder blades. A suspension of mercaptosuccinic acid (MSA, 400 mg/kg, Sigma-Aldrich) in 40% Pluronic hydrogel was prepared and 25 μl administered topically into the wound. Wounds were dressed with a sterile Tegaderm dressing (3M, Bracknell, UK) which prevented wound contamination, maintained a moist environment and ensured animals would not clean their wounds. Bepanthen Ointment (Bayer, Reading, UK) was applied to the uncovered, depilated skin surrounding the dressing to minimise any irritation.

Mice were individually housed in conventional cages in a 28C warm box throughout the experiment, with additional bedding, which effectively enabled these elderly mice to maintain their body weight. Animals were provided with ‘mash’ (standard chow rehydrated in sterile water) throughout the protocol and were monitored as outlined in Fig. S3. At Day 7 post-wounding, CWs were carefully debrided to remove dead and sloughing tissue. Briefly, mice were anaesthetized with isoflurane by inhalation, placed on a heated mat and administered Buprenorphine (0.05 mg/kg sc). The existing Tegaderm dressing was carefully removed, ensuring the wound was not pulled in the process. Debridement was performed using 4.25 inch dissection scissors and curved tweezers (Sigma-Aldrich) Vehicle (PBS) or BMS-470539 (MC1R-Ag) were delivered topically by pipette into the wound cavity immediately after debridement (100 μl in a 30% Pluronic F-127 gel; Sigma Aldrich). Wound areas were calculated using ImageJ software.

### Evans Blue assay of lymphatic drainage

Evans blue dye (Sigma-Aldrich, Poole) was dissolved in sterile saline solution to a concentration of 0.5% w/v and 10 μl was injected into the cutaneous wound bed 8 hours prior to cull. The gross appearance of the tissue was recorded using a digital camera immediately following cull. The skin draining lymph nodes (axillary, brachial and inguinal) were then harvested into 200 μl of formamide and incubated for 24 h at 55°C to extract the dye. The tissue pieces were discarded, debris removed via centrifugation and absorbance read in triplicate at 610 nm.

### Wound processing

Acute wounds were harvested on days 1, 3, 7, 10 and 14 post-wounding, fixed in 4% PFA (overnight at 4C on a rocker, Sigma Aldrich), washed 3 x 5 min in PBS, transferred to 70% ethanol and then embedded in paraffin. Day 1 and 3 wounds were cut in half using a scalpel and sections (10 µm) taken from the middle of the wound. Day 7, 10 and 14 wounds were sectioned through the wound beyond the midpoint and wound centres identified by staining with Haematoxylin (Gills No.3) and Eosin (Sigma). All CWs were bisected.

### Histology

FFPE sections were deparaffinised and rehydrated in: xylene (2 x 3 min), 100% ethanol (2 x 3 nub), 90% ethanol (3 min) and 70% ethanol (3 min). Sections were placed in running tap water before being submerged in haematoxylin (Sigma-Aldrich, Dorset, UK) for 5 s – 1 min depending on staining intensity desired. Excess haematoxylin was rinsed off in tap water twice. Slides were then briefly submerged in acid alcohol (300 ml dH20, 700 ml 100% ethanol, 3 ml concentrated HCL; Sigma-Aldrich), followed by tap water wash. Sections were submerged in eosin (Sigma-Aldrich) for 30 s to provide a counterstain to the haematoxylin. After tap water washes (x 3), sections were dehydrated for mounting in sequential baths of 70%, 90%, 100% ethanol and xylene (x 2) for 30 seconds each. DPX mountant (Sigma-Aldrich) was applied to the sections, followed by a Menzel microscope coverslip (25 x 60 mm, 0.13 mm thick; Thermofisher Scientific, Renfrew, UK). For Picrosirius Red (PSR) staining for collagen, the above protocol was followed but with PSR (0.5 g Sirius Red, 1.3% picric acid in dH2O; Sigma-Aldrich) for 1 hour and no eosin counterstain. For Prussian Blue staining, the above protocol was followed but with Prussian Blue (10% HCl, 5% potassium ferrocyanide in dH2O used rather than haematoxylin.

### Immunohistochemistry

Skin sections were deparaffinised and rehydrated as described above. Sections were encircled with a hydrophobic barrier using an ImmEdge pen (Vector labs, Peterborough, UK). Antigen retrieval was carried out using Proteinase K (5 μg/ml, 7 mins, Dako, UK). Sections were rinsed in running tap water for 5 min, blocked (1% BSA, 0.1% Gelatin, 0.5% Triton X-100; Sigma-Aldrich) for 30 min at room temperature. Sections were incubated in primary antibody diluted in block, overnight at 4C in a humidified chamber. After PBS-0.1% Tween-20 wash, an ImmPRESS polymer detection kit was used, followed by a DAB kit, as per the manufacturer’s instructions (Vector labs). Sections were counterstained with haematoxylin, dehydrated and mounted as described above.

### Microscopy

Tissue sections were imaged on an Axioscan Z1 slide scanner (Zeiss, Cambridge, UK) using the 20x air objective. Immunofluorescence images were taken using a Leica DM2500 microscope, Leica DFC7000 camera and LAS software (Leica Microsystems, GmbH, Milton Keynes, UK), using the 20 x air objective. Numbers of leukocytes per field of view (FOV) were calculated using thresholded 32-bit images and the “analyze particles” function in FIJI software. Blood and lymphatic vessels were quantified by thresholding on fluorescent area in FIJI software.

### Collagen alignment

Collagen fibre orientation was desired to be able to identify the level to which wounds recovered their natural structural components during the wound healing process. To undertake this, we developed a MATLAB code which first isolated collagen fibrils from non-fibrous tissue using a colour-based segmentation algorithm. First, the user selected image is converted from an RGB image to a two-dimensional matrix of type *m* × *n* × 3. Where *m* and *n* represent the row and column indices respectively. Our code (which we name FIBRAL) converts the image colour space from RGB to L*a*b to more accurately predict small colour differences, key for PSR staining in which small variation detection is required (31). Both the full a-channel and positive range of the b-channel were isolated and superimposed. This new matrix was transformed into grayscale with contrast and brightness increased by 20% and 5% respectively.

To determine the spatial distribution of collagen fibres, a two-dimensional Fourier transformation was performed on the image. This data was transformed from cartesian to polar coordinates by grouping predominant frequency-component orientations of the fibres in 1-degree increments from 0 to 180 degrees. The total intensity at each angle was calculated by summing the grayscale intensity values of each pixel contained within the group. Large intensity values represented the summation of multiple sinusoidal waves with large amplitude signifying major fibre alignment along the respective orientation. From this data, a comparative analysis can be conducted whereby the dot product was calculated using each permutation of resulting vectors (parallel vectors would provide a dot product of 1, with orthogonal vectors presenting a dot product value of -1). All results were normalised to lie between 0 and 1 with a directionality quotient of 1 representing a parallel field of fibres and a directionality quotient of 0 representing full fibre orientation across all angles in the tissue. The code was validated against images with dominant fibre orientations and combinations thereof. Full details of the coding approach are included in the supplementary material.

### Wound Optical Coherence Tomography Angiography

OCTA imaging is a non-invasive imaging method which utilises light to produce cross-sectional images of tissue architecture *in situ* and in real time. This form of imaging allowed the tracking of wound healing parameters such as reepithelialisation within a single mouse over a time course of healing. Additionally OCTA provides information on the perfusion of new blood vessels which cannot be obtained utilising other methods such as immunofluorescence staining.

Equipment used in this study were designed and developed by Professor Zhihong Huang, Dr. Kanheng Zhou, Mr. Yubo Ji and Mr. Duo Zhang (University of Dundee (32,33)). Mice were anaesthetised at the indicated time points, placed on a heatpad and ultrasound transmission gel (Anagel, Ana Wiz Ltd, Surrey, UK) was applied between the OCT probe and wound to enhance refractive index coupling. A clinical prototype handheld OCTA system was utilised to image dorsal skin wounds. A 200-kHz vertical-cavity surface-emitting swept source laser (5 mW) (SL132120, Thorlabs Inc., Newton, USA) was used as the light source in the system paired with a XY galvo-scanner (6210H, Cambridge technology, Bedford, USA) and a 5X objective lens (LSM03, Thorlabs Inc., Newton, USA). With this configuration, the handheld probe provided a scanning range of 10 × 10 mm2. To generate high resolution 3D OCTA images for skin architecture and vasculature, the following B-M scanning protocol was applied: Firstly, A-lines were acquired consecutively with a 10 μm interval along the fast scanning axis to generate a B-frame. Each B-frame was then repeated four times at the same location for blood flow extraction. Next, the described repeated B-scans were looped consecutively along the slow scanning axis also with a 10 μm interval to form a complete 3D volume scan. The final volumetric dataset comprised 600 A-lines and 600 cross-sectional planes (2400 B-frames). Data was collected through a customised LabVIEW (NI, Texas, USA) interface for processing.

### HMEC-1 culture and transfection

HMEC-1 cells (ATCC: CRL-3243) were cultured in EGM-2 MV (EBM-2 added with growth factors and other supplements, PromoCell). Lipofectamine RNAiMAX (Thermo Fisher Scientific) was used to transfect HMEC-1 with siRNA control or siRNA MC1R oligonucleotides (Qiagen) (20 nM final concentration).

### Endothelial Tube Formation Assay

Matrigel (50 μL/well) (BD Matrigel Basement Membrane Matrix, BD Biosciences) was added to the wells in a 96-well plate and allowed to polymerize at 37 °C for 30 min. HMEC-1 cells, an immortalised human microvascular endothelial cell line (34) (15,000 cells) previously transfected with siRNA-control or siRNA MC1R were added to the top of the Matrigel and treated with MC1R agonist BMS-470539. After incubation for 8 hr, tube formation was assessed by acquiring four random microscopic fields (magnification ×10) by using Nikon Eclipse Ti inverted microscope. Total tube length and branching points were assessed using Angiogenesis Analyzer for ImageJ analysis software (35). For confocal imaging, cells were fixed (4% PFA, 15 m), permeablised (0.1% Triton X-100, 15 min), blocked (1% BSA, 0.1% Gelatin, 0.5%, 30 min) and stained according to manufacturer’s instructions with Phalloidin Alexa 647 (ThermoFisher Scientific) and DAPI (Sigma Aldrich).

### Human Tissue

FFPE sections of human CWs were provided by NHS Lothian Tissue Bank (ethical approval: 15/ED/0094) with approval from the Tissue Governance Committee (SR612, SR1368, SR736). Inclusion criteria were pressure ulcers, diabetic foot ulcers and venous leg ulcers from male and female donors. Exclusion criteria were patients with cancer and pressure ulcers from patients with diabetes. Prior to receipt, all samples were anonymised.

### Data analysis of GSE165816

Preprocessed and quality filtered scRNA-seq data from Theocharidis G *et al* was downloaded and imported into R (v4.2.2) for reanalysis (36). Count data was normalized using the SCTransform algorithm in the Seurat Bioconductor package (v4.4) that uses regularized negative binomial models for normalizing sparse single-cell data. The normalized expression profiles of the samples were merged, before unsupervised and supervised analysis using various R and Bioconductor packages. The quality filtering on scRNASeq data used multiple filtering parameters to remove cells with >10**%** of mitochondrial genes, cells expressing <200 genes and genes only uniquely expressed in <3 cells. Unsupervised principal component analysis (PCA) was performed on variable genes to identify principal components, which captured the most variance across the samples. These principal components were used as an input for Uniform Manifold Approximation and Projection (UMAP) analysis to determine and visualise the overall relationship among the cells. Cell clusters with similar transcriptome profiles were identified by shared nearest neighbour (SNN) analysis; clusters were subject to marker detection and subsequently annotated to different cell types based on the expression of specific well-established cell marker transcripts as per annotation in Theocharidis G *et al*.

### POMC-MC1R Expression and Systematic inference of cell-cell communication

In order to handle low abundance transcripts POMC and MC1R, expression data within cell type clusters were pooled, generating normalized transcript per million (nTPM) for each gene and cell type. ‘RC’ (relative count) with a scale factor of 1e6 was used for the Seurat normalisation method. CellChat was used to systematically infer cell-cell communication by integrating scRNA-seq data with prior ligand-receptor interaction database CellChatDB (Jin et al., 2021). CellChat prediction of putative cell-cell communications occur in three steps. First, CellChat identifies differentially over-expressed ligands and receptors for each cell group.

Second, CellChat quantifies the communication probability between two interacting cell groups based on the average expression values of a ligand by one cell group and that of a receptor by another cell group, as well as their cofactors. With the low expression level of POMC-MC1R, "truncatedMean" function with a trim of 0.01 was exploited to calculate the communication probabilities. Third, CellChat infers significant interactions using permutation tests. CellChat outputs an intercellular communication network for each ligand-receptor pair, where the calculated communication probabilities are assigned as edge weights to quantify the interaction strength. An intercellular communication network of a signaling pathway computed by summarizing the communication probabilities of POMC-MC1R ligand-receptor pairs.

## Statistical analysis

Student’s *t*-test, one-way and two-way ANOVA with Bonferroni’s multiple comparison test were performed using GraphPad Prism 9.0 software and detailed in the respective Figure legends.

## Diagrams

Biorender was used to create the schematics in Figures 1, 3 and 4.

## Data availability statement

Data will be deposited in the open access repository Figshare prior to publication. The acute and chronic wound protocols are in preparation for publication as a methods paper and will be made available upon request. Data comprising Fig 5A-D were derived from analysis of GSE165816 (36).

## Acknowledgments

We acknowledge the support of the QMRI SURF facilities for histological services and BVS (Bioresearch and Veterinary Services), especially Dr Nacho Vinuela-Fernandez for assistance with animal studies. We thank Professor Ian Jackson for providing the MC1Re/e mouse line and Dr Andrew Tambyraja and Professor Keith Harding for helpful insights on CWs from a clinical perspective and Professor Paul Martin for general discussions. We thank the Wolfson Bioimaging Facility for assistance with Incucyte experiments and Dr Matthieu Vermeren (CALM services) for microscopy input. This work has made use of the resources provided by the Edinburgh Compute and Data Facility (ECDF). Thanks also to Prof David Dockrell and Prof Julia Dorin for mentorship. This work was funded by a Sir Henry Dale Fellowship (202581/Z/16/Z), Elizabeth Blackwell Early Career Fellowship (Wellcome ISSF) and a University of Edinburgh Chancellors Fellowship awarded to JC. HR was supported with a Chancellors Fellowship PhD Studentship. Work in MC lab was funded by an EPSRC New Investigator grant EP/S019847/1 and CB is funded by a Carnegie Trust for Scotland PhD studentship.

## Contributions

YN, HR and SAW performed experiments, analysed the data and wrote the manuscript. GK and AP performed experiments and analysed the data. KZ and YJ performed OCT imaging and analysis under the supervision of ZH. CB, YC and MC developed FIBRAL. AP performed experiments and analysed data. AMK assisted with scRNAseq data analysis under the supervision of SJF. AB provided expert advice as an inflammatory dermatopathologist. AC advised on the design and execution of *in vitro* angiogenesis experiments, performed and analysed these experiments. JLC conceived and planned the project, performed the experiments, analysed and interpreted the data, wrote the manuscript, obtained funding, and supervised the project.

