## Supplemental Figures 1-4 for "MC1R reduces scarring and rescues stalled healing in a preclinical chronic wound model"

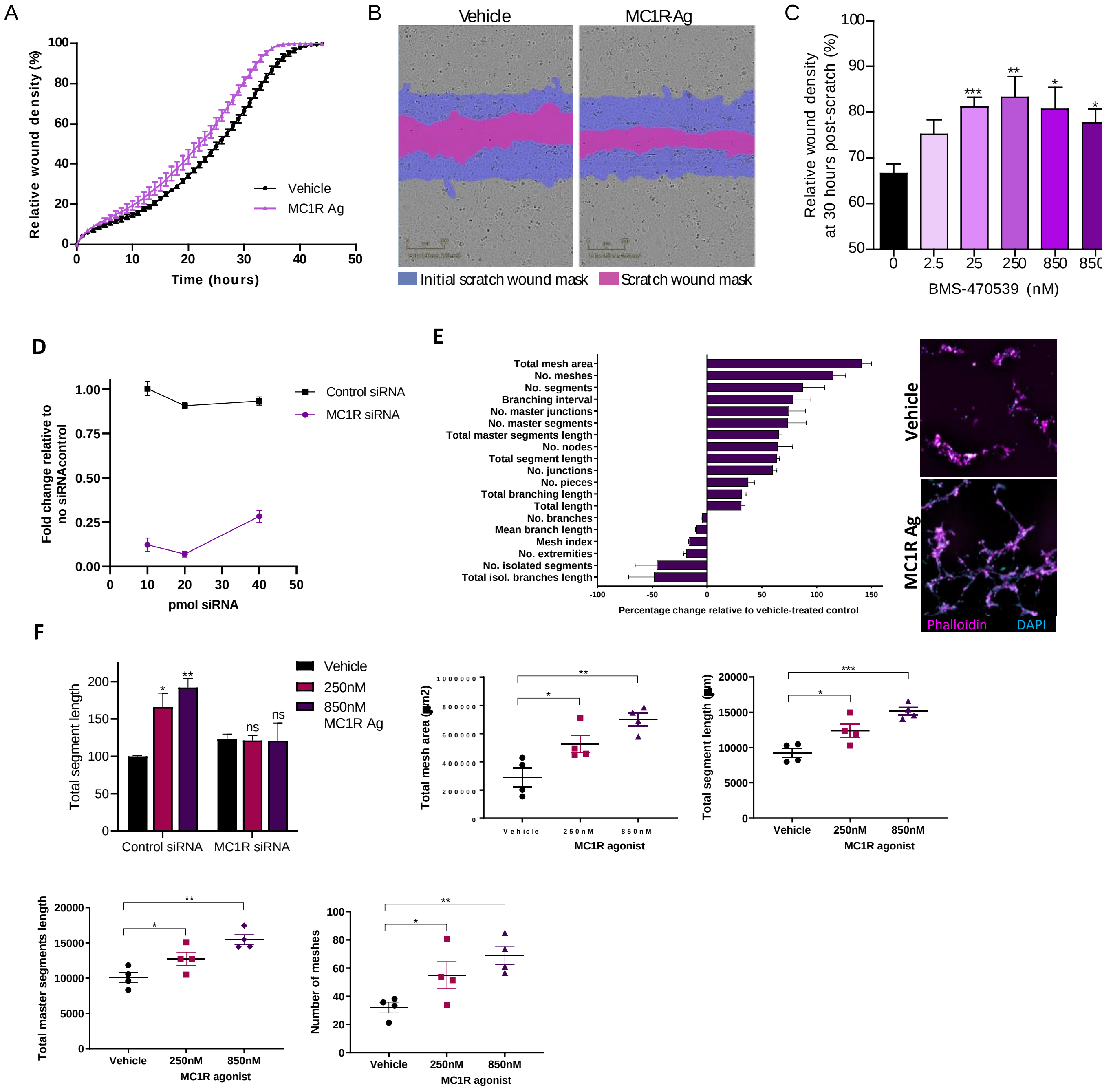


**Supplementary Figure 1 – BMS-470539 directly drives *in vitro* scratch wound closure and angiogenesis through MC1R**

**(A-C)** HaCaTs (human keratinocytes) were allowed to migrate for 48 hr to fill in a “scratch wound” *in vitro* in the presence of BMS-470539 or media control. **(A)** Time course of scratch wound closure with vehicle or MC1R-Ag treatment (250 nM BMS-470539). **(B)** Representative images taken at 7 h Dose response of scratch wound closure. **(C)** Dose response of BMS-470539–induced scratch wound closure. **(D)** Optimisation of MC1R siRNA knockdown. A dose of 20 pM was chosen for subsequent experiments. **(E)** Percentage change in *in vitro* angiogenesis parameters with MC1R-Ag-treatment (850 nM BMS-470539) relative to vehicle-treated control (four independent experiment). Representative images of Matrigel angiogenesis assay with HMEC-1 cells treated with vehicle or MC1R agonist BMS-470539 (250 nM). Cells stained with Phalloidin (magenta) and DAPI (blue). **(F)** Quantification of four key *in vitro* angiogenesis parameters in the presence and absence of MC1R agonist (250 nM and 850 nM). Data are expressed as mean ± SEM. Four independent experiments, each with four technical replicates. Data are expressed as mean ± SEM. * $p<0.05$, ** $p<0.01$, *** $p<0.001$ by one-way ANOVA with Bonferroni’s multiple comparison test (A) relative to vehicle-treated wounds.

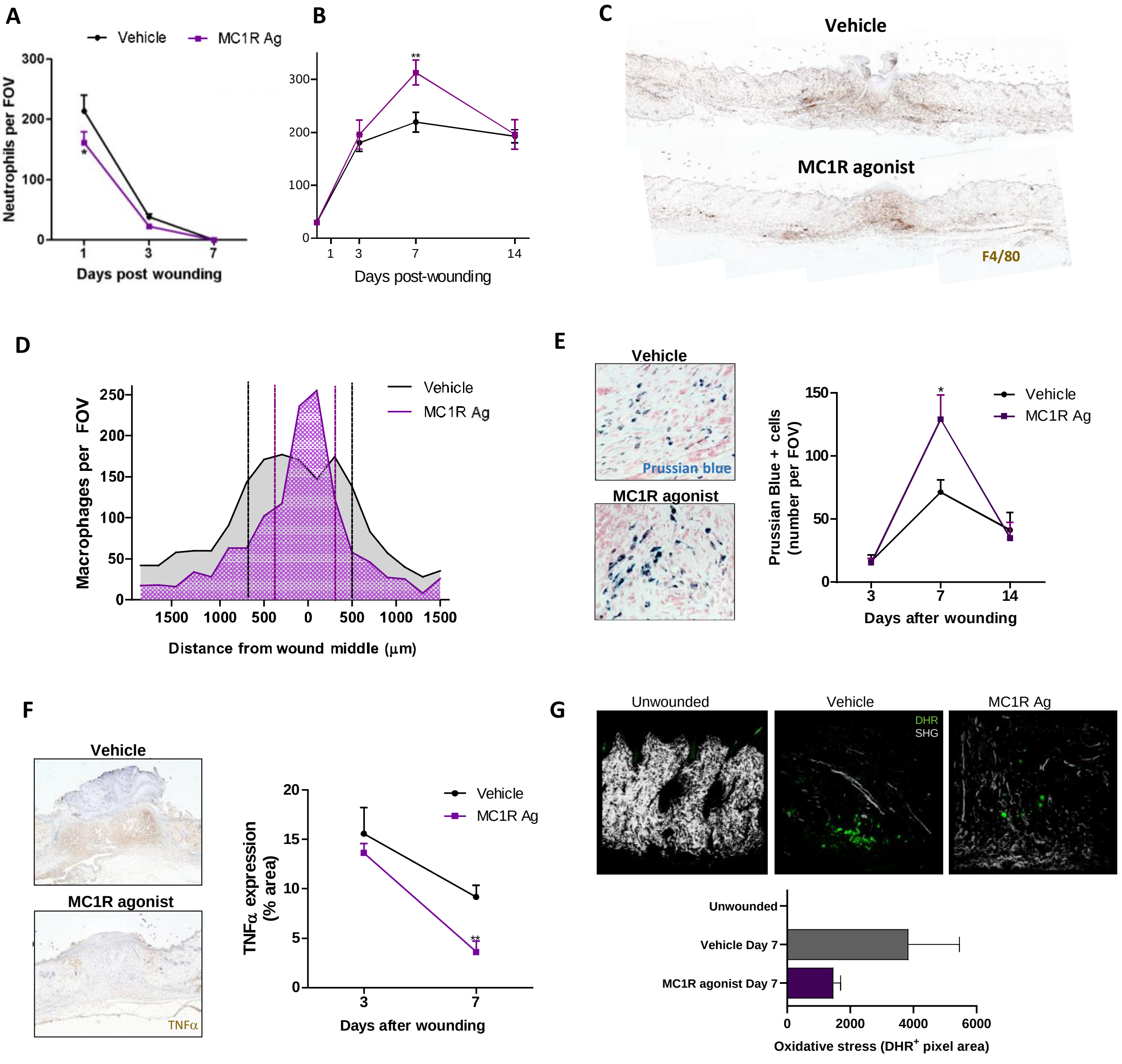


**Supplementary Figure 2 – MC1R agonist modulates the wound microenvironment**

**(A)** Quantification of wound bed neutrophil recruitment (Ly6G$^{+}$ cell) 1-7 dpi. **(B)** Quantification of wound bed monocyte-macrophage recruitment (F4/80$^{+}$ cell) 3-14 dpi. **(C)** Macrophage immunostaining (F4/80) through the mid-point of wounds 7 dpi. **(D)** Quantification of macrophage distribution across wounds and adjacent tissue. Wound margins denoted by solid black (vehicle) and purple (MC1R-Ag) lines. **(E)** Representative hpf and quantification of Prussian blue$^{+}$ iron-loaded cells within the wound 3-14 dpi. Images from 7 dpi sections with Eosin counterstain. **(F)** Quantification and representative lpf (5x objective) of TNFα expression (brown) with haematoxylin counterstain at 7 dpi. **(G)** Representative images and quantification of oxidative stress (dihydrorhodamine$^{+}$ pixel area; green). Second harmonic generation (SHG; collagen, white). Data are expressed as mean ± SEM. * $p<0.05$, ** $p<0.01$, *** $p<0.001$ by two-way ANOVA with Bonferroni's multiple comparison test (A, B, E, F) relative to vehicle-treated wounds. Seven to eleven wounds per group from three independent experiments. FOV, field of view.

**A**

Ethical approval number

Animal number

| dpi | Date | Wound | | Wound Area | | | | Activity and Movement | Behaviour | | | Irritation | Weight | Weight (%) | Pain | Bepanthen applied | Hair regrowth | Dressing | | Photo Taken | Mash | Notes | Monitor (initials) | NVS spoken to | 20% weight threshold | 10% weight threshold |
|---|---|---|---|---|---|---|---|---|---|---|---|---|---|---|---|---|---|---|---|---|---|---|---|---|---|---|
| | | Score | Exudate | Score | Size | Appearance | | Score | Score | Scratching | Shaking | | | | | | | Appearance | Changes | | | | | | | |
| | | | | | | Dead white tissue | Red ring around tissue | | | | | | | | | | | | | | | | | | | |
| BASELINE | 10.3.22 | | | | | | | | | | | | 36 | | | | | | | | Yes | | | | 28.8 | 32.4 |
| 1 | 11.3.22 | 1 | No Exudate | 1 | Growing larger | Yes | No | 0 | 0 | No | No | No | 35.8 | 99.44% | 0 | No | No | Adherent | No | | Yes | | | | | |
| 2 | 12.3.22 | 1 | Clear Exudate | 1 | No Change | Yes | Yes | 0 | 0 | No | No | No | 34.2 | 95.00% | 0 | No | No | Adherent | No | | Yes | | | | | |
| 3 | 13.3.22 | 1 | Clear Exudate | 1 | No Change | Yes | Yes | 0 | 0 | No | No | No | 35.6 | 98.89% | 0 | No | No | Adherent | No | | Yes | bit of crustiness next to dressing (leaked wound exudate) | | | | |
| 4 | 14.3.22 | 1 | Clear Exudate | 1 | No Change | Yes | Yes | 0 | 0 | No | No | No | 36.5 | 101.39% | 0 | No | No | Adherent | No | | Yes | bit of crustiness next to dressing (leaked wound exudate) | | | | |
| 5 | 15.3.22 | 1 | Clear Exudate | 1 | No Change | Yes | Yes | 0 | 0 | Yes | No | No | 36.2 | 100.56% | 0 | No | No | Adherent | No | | Yes | bit of crustiness next to dressing (leaked wound exudate) | | | | |
| 6 | 16.3.22 | 1 | Clear Exudate | 1 | No Change | Yes | No | 0 | 0 | No | No | No | 34.2 | 95.00% | 0 | No | No | Edges lifted | No | | Yes | | | | | |
| 7 | 17.3.22 | 1 | Clear Exudate | 1 | No Change | Yes | No | 0 | 0 | No | No | No | 34.2 | 95.00% | 0 | No | No | Adherent | Yes | | Yes | Debrided, dressing change + PL treatment | | | | |
| 8 | 18.3.22 | 1 | Clear Exudate | 1 | No Change | | | 0 | 0 | No | No | No | 34.8 | 96.67% | 0 | No | No | Adherent | No | | Yes | | | | | |
| 9 | 19.3.22 | 1 | Clear Exudate | 1 | No Change | | | 0 | 0 | Yes | No | No | 34.3 | 95.28% | 0 | No | No | Adherent | | | Yes | bit of crustiness next to dressing (leaked wound exudate) | | | | |
| 10 | 20.3.22 | 1 | Clear Exudate | 1 | No Change | | | 0 | 0 | No | No | No | 34.2 | 95.00% | 0 | No | No | Adherent | Yes | | Yes | Dressing change and PL treatment. | | | | |
| 11 | 21.3.22 | 1 | Clear Exudate | 1 | No Change | | | 0 | 0 | No | No | No | 33.7 | 93.61% | 0 | No | No | Adherent | No | | Yes | | | | | |
| 12 | 22.3.22 | 1 | Clear Exudate | 1 | No Change | | | 0 | 0 | No | No | No | 33.6 | 93.33% | 0 | No | No | Edges lifted | Yes | | Yes | Large, sloughy, wet wound. Crusty. PL and dressing change. | | | | |
| 13 | 23.3.22 | 1 | Clear Exudate | 1 | No Change | | | 0 | 0 | No | No | No | 33.9 | 94.17% | 0 | No | No | Adherent | No | | Yes | bit of crustiness next to dressing (leaked wound exudate) | | | | |
| 14 | 24.3.22 | 1 | Clear Exudate | 1 | No Change | | | 0 | 0 | No | No | No | 34 | 94.44% | 0 | No | No | Adherent | No | | Yes | | | | | |

**B**

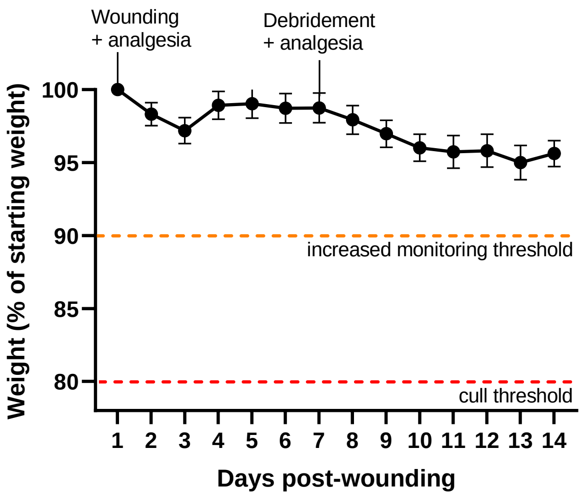


**C**

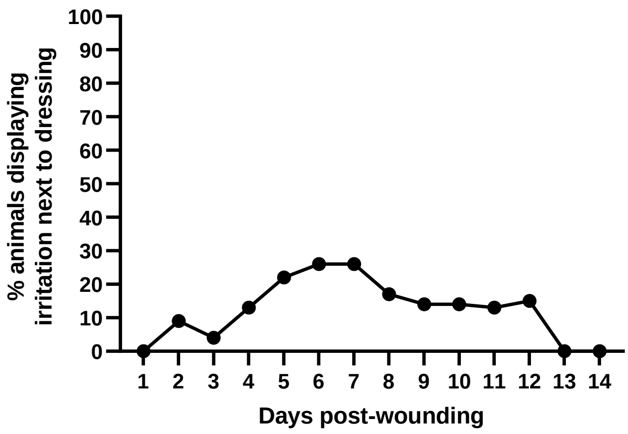


**Supplementary Figure 3 – Murine chronic wounds are humane and very well tolerated**

**(A)** Example monitoring record of wound features, animal behaviour and weight to ensure humane and ethical treatment. **(B)** Minimal weight loss indicates that the chronic wound protocol is well tolerated by the aged mice. Data displayed as mean ± SEM (n=24). **(C)** Up to 30% of mice displayed minor skin irritation next to the Tegaderm dressing. When this was observed Bepanthen cream was applied to soothe the area and prevent the irritation from worsening (n=24).

| | Patient details | | | | |
|---|---|---|---|---|---|
| Code | Sex | Age | Wound location | Duration of wound | Diagnosis |
| 1D | F | 77 | Leg | 2 years | Diabetic ulcer |
| 2D | M | 59 | Ankle | 7 years | Diabetic ulcer |
| 4D | F | 59 | left hallux | 4 years | Diabetic ulcer |
| 5D | F | 60 | Left foot, 3rd toe | 5 years | Diabetic ulcer |
| 8D | M | 60 | right shin | 1 year | Diabetic ulcer |
| 9D | F | 57 | lateral anterio left leg | 1 year | Diabetic ulcer |
| 10D | F | 73 | left posterior heel | 6 months | Diabetic ulcer |
| 11D | F | 59 | top of right middle finger | 2 weeks | Diabetic ulcer |
| 14D | M | 87 | lateral border of left foot | 3 months | Diabetic ulcer |
| 4P | F | 44 | Buttock | 6 months | Pressure ulcer |
| 8P | F | 73 | right medial heel | 8 months | Pressure ulcer |
| 9P | F | 78 | right greater trochanter | not recorded | Pressure ulcer |
| 12P | F | 68 | left ischium | not recorded | Pressure ulcer |
| 13P | M | 72 | left ischium | 7 months | Pressure ulcer |
| 17P | F | 52 | left shoulder | 2 months | Pressure ulcer |
| 18P | M | 12 | sacral area | 1 year | Pressure ulcer |
| 6V | F | 83 | left ankle | 6 months | Venous ulcer |
| 7V | F | 77 | right mid anterior shin | 1 year | Venous ulcer |
| 8V | M | 52 | left medial malleolus | 10 months | Venous ulcer |
| 9V | M | 89 | left ankle | 5 months | Venous ulcer |
| 10V | M | 62 | left lower leg | 6 weeks | Venous ulcer |
| 11V | F | 62 | left medial malleolus | 2 weeeks | Venous ulcer |
| 13V | F | 71 | lateral aspect right foot | 9 months | Venous ulcer |
| 15V | F | 77 | left medial malleolus | 6 months | Venous ulcer |
| 16V | M | 67 | right leg | 1 year | Venous ulcer |
| 17V | M | 69 | back of lower right calf | 3 years | Venous ulcer |
| 18V | F | 89 | left shin | 3 months | Venous ulcer |
| 19V | M | 67 | lower left leg | 1 year | Venous ulcer |
| 20V | F | 69 | right medial malleolus | 6 years | Venous ulcer |
| 21V | M | 94 | left anterior fibula | 4 months | Venous ulcer |
| 22V | M | 34 | lateral left lower leg | 6 months | Venous ulcer |
| 28V | M | 63 | left ankle | 1 year | Venous ulcer |
| 31V | F | 73 | left ankle | 2 years | Venous ulcer |
| 32V | F | 88 | right posterior distal leg | unknown | Venous ulcer |
| 33V | F | 87 | lower left leg | 3 years | Venous ulcer |
| 02_VL | M | 59 | anteromedial portion of left shin | 1 year | Venous ulcer |
| 04_VL | F | 74 | right lower leg | 7 months | Venous ulcer |
| 05_VL | M | 80 | right lateral malleolus | unknown | Venous ulcer |
| 06_VL | F | 82 | gaiter region of left leg | 3 years | Venous ulcer |
| 07_VL | F | 86 | right medial malleolus | 7 months | Venous ulcer |
| 08_VL | F | 71 | left medial ankle | 5 months | Venous ulcer |
| 09_VL | M | 92 | right lower medial leg | 8 weeks | Venous ulcer |
| 10_VL | F | 81 | Left lower leg | 2 months | Venous ulcer |

**Table 1 – Donor information**
